# Forgetting Enhances Episodic Control with Structured Memories

**DOI:** 10.1101/2021.08.11.455968

**Authors:** Annik Yalnizyan-Carson, Blake A. Richards

## Abstract

Forgetting is a normal process in healthy brains, and evidence suggests that the mammalian brain forgets more than is required based on limitations of mnemonic capacity. Episodic memories, in particular, are liable to be forgotten over time. Researchers have hypothesized that it may be beneficial for decision making to forget episodic memories over time. Reinforcement learning offers a normative framework in which to test such hypotheses. Here, we show that a reinforcement learning agent that uses an episodic memory cache to find rewards in maze environments can forget a large percentage of older memories without any performance impairments, if they utilize mnemonic representations that contain structural information about space. Moreover, we show that some forgetting can actually provide a benefit in performance compared to agents with unbounded memories. Our analyses of the agents show that forgetting reduces the influence of outdated information and states which are not frequently visited on the policies produced by the episodic control system. These results support the hypothesis that some degree of forgetting can be beneficial for decision making, which can help to explain why the brain forgets more than is required by capacity limitations.

## 1 INTRODUCTION

Many people bemoan their tendency to forget, and assume that if it was possible, it would be desirable to remember everything that had ever happened to them. Yet, evidence from psychology and neuroscience suggests that the mammalian brain has the capacity to store far more episodic memories than it does, and that healthy brains actually engage in active forgetting of episodic memories. For example, some individuals with a syndrome known as Highly Superior Autobiographical Memory (HSAM), are capable of remembering almost everything that has ever happened to them (Parker et al., 2006; Leport et al., 2016), but these individuals assert that this is a detriment for them, not an advantage. As well, neurobiological studies of forgetting have shown that a diverse array of molecular and cellular mechanisms promote the active forgetting of information (Akers et al. (2014); Epp et al. (2016); Shuai et al. (2010); Migues et al. (2016); Berry et al. (2012) – see also Wixted (2004); Hardt et al. (2013); Richards and Frankland (2017); Anderson and Hulbert (2021) for reviews). In fact, researchers can prevent forgetting in animal models by interfering with these mechanisms, demonstrating that in principle, animals could remember more than they do (Berry et al., 2012; Shuai et al., 2010; Akers et al., 2014).

Why would our brains actively forget what has happened to us? It has been hypothesized that transient memories may provide a better substrate for decision making, as they would render animals more flexible and better at generalization (Mosha and Robertson, 2016; Robertson, 2018; Hardt et al., 2013; Richards and Frankland, 2017). This “beneficial forgetting” hypothesis is supported by some animal studies demonstrating that artificially reducing forgetting can impair reversal learning and reduce generalization of learned associations (Epp et al., 2016; Shuai et al., 2010; Migues et al., 2016). For example, reducing AMPA receptor internalization in the hippocampus prevents the generalization of contextual fear memories (Migues et al., 2016).

However, it is difficult to fully examine the validity of this normative hypothesis in real-life experiments. In the previous example (Migues et al., 2016), it is difficult to say exactly why reduced AMPA receptor internalization prevents generalization—is it due to reduced forgetting, or some other downstream affects of the experimental manipulation? Modelling and simulation provide a means of exploring the computational validity of the beneficial forgetting hypothesis in a fully controlled manner (Brea et al., 2014; Murre et al., 2013; Toyama et al., 2019). In particular, reinforcement learning (RL) from artificial intelligence (AI) provides a normative framework that is ideal for understanding the role of memory in decision making (Niv, 2009; Gershman and Daw, 2017; Dolan and Dayan, 2013). In particular, episodic control, an approach in RL that utilizes one-shot memories of past events to shape an agent’s policy (Lengyel and Dayan, 2007; Pritzel et al., 2017; Blundell et al., 2016; Ritter et al., 2018), is ideal for exploring the potential impact of forgetting on decision making and computation.

Therefore, to determine the validity of the beneficial forgetting hypothesis, we used an episodic control agent trained to forage for rewards in a series of maze environments. We manipulated both the underlying representations used for memory storage and the degree to which the episodic memory cache forgot old information. We find that when memories are stored using structured representations, moderate amounts of forgetting will not only leave foraging abilities intact, but will actually produce some modest performance improvements. We find that these performance gains result from the fact that forgetting with structured mnemonic representations eliminates outdated and noisy information from the memory cache. As a result, the agent’s episodic control system will produce policies that are more consistent over local neighbourhoods of state space. As well, forgetting with structured representations can preserve the confidence of the agent’s policies, particularly those near the goal. Altogether, these results support the beneficial forgetting hypothesis. They show that if an agent is using records of past experiences to guide their actions then moderate amounts of forgetting can help to produce more consistent decisions that generalize across space.

## 2 MATERIALS AND METHODS

### 2.1 Reinforcement Learning Formalization

Our episodic control model was designed to solve a reinforcement learning task. The reinforcement learning problem is described as a Markov decision process (MDP) for an “agent” that must decide on actions that will maximize long-term performance. An MDP is composed of a set of discrete states (*s* ∈ 𝒮) sampled over time, *t*, a set of discrete actions (*a* ∈ 𝒜), a state-transition probability distribution *P* (*s*′|*s*_*t*_ = *s, a*_*t*_ = *a*), a reward function *R*(*s*) = *r*, and a discount factor *γ* ∈ [0, 1] to weigh the relative value of current versus future rewards (we set *γ*=0.98 for all simulations). The transition distribution *P* (*s*_*t*+1_ = *s*′|*s*_*t*_ = *s, a*_*t*_ = *a*) specifies the probability of transitioning from state *s* to a successor state *s*′ when taking action *a*. The reward function *R*(*s*) specifies the reward given in state *s*. The transition distribution and reward function describe the statistics of the environment, but are not explicitly available to the agent; the agent only observes samples of state-action-reward-next state transitions (*s*_*t*_ = *s, a*_*t*_ = *a, r*_*t*_ = *r, s*_*t*+1_ = *s*′). The policy *π*(*a* | *s*_*t*_ = *s*) specifies a probability distribution over available actions, from which the agent samples in order to select an action in the environment. The return at time *t, G*_*t*_, is the temporally discounted sum of rewards from the present time-step into the future:

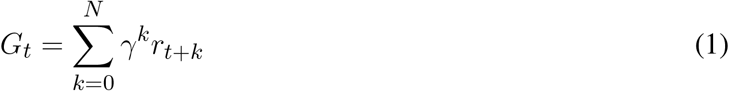

The aim of the reinforcement learning task is for the agent to generate policies which will allow it to maximize the return.

### 2.2 Environments and Foraging Task Design

Simulations of the foraging task were carried out in four different grid-world environments: (1) open field, (2) separated field, (3) four rooms, and (4) tunnel (Fig. 1A). Each environment was designed as a 20×20 grid of states connected along a square lattice, with four possible actions Down, Up, Left, and Right in each state. In three of the four test environments (separated field, four rooms, tunnel), obstacles were present in some states. These obstacle states were removed from the graph such that there were no edges connecting obstacles to other states. That is, the graph adjacency matrix was updated such that *A*(*s, o*) = 0 for any state *s* and adjacent obstacle state *o*. Actions which would result in the agent moving into a barrier or boundary returned the agent’s current state.

**Figure 1.**
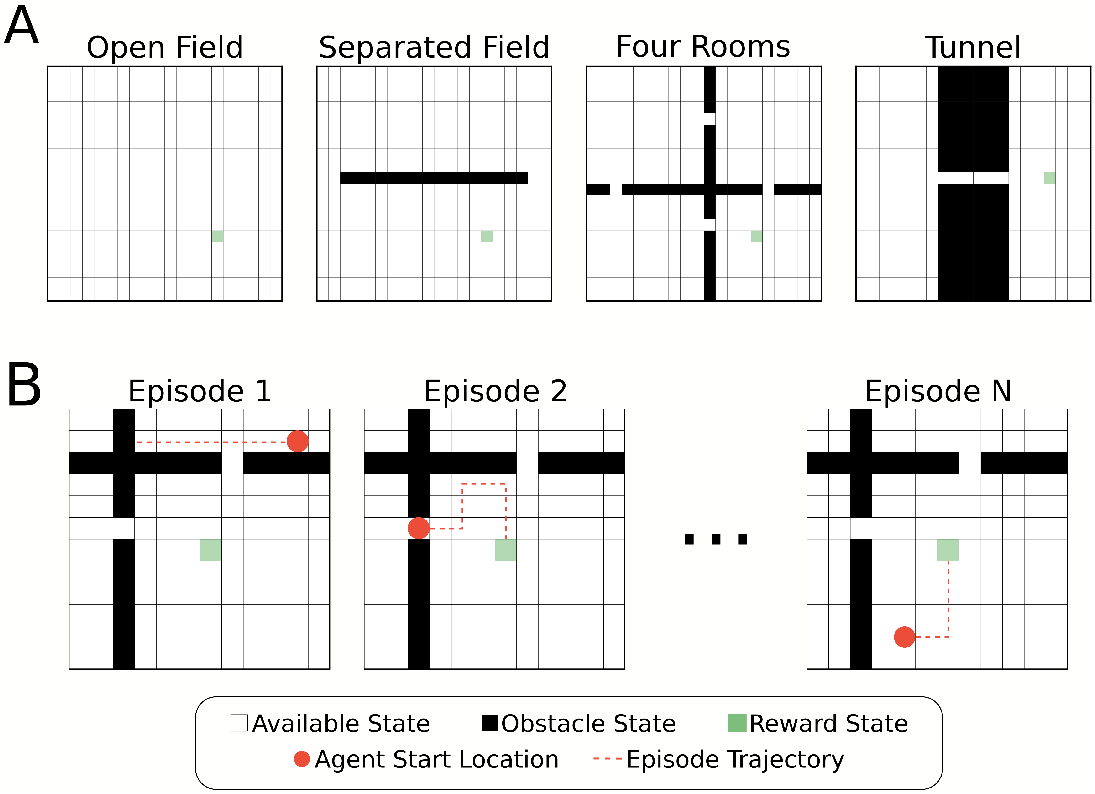
Illustration of the environments and foraging task. (A) Four different grid-world environments were used for the foraging task to test the effects of memory restriction on episodic control. The environments differed in placement of obstacle states. Partitioning the plane created bottleneck states through which the agent must successfully navigate to reach the reward location. (B) Example episodes from the foraging tasks. The agent’s starting location at the beginning of each episode is chosen at random from the available states in the environment.

Each environment contained a single location associated with a reward state, *s*\*, (*R*(*s*\*) = 10) that the agent had to “forage” for. All other states were associated with a small penalization (*R*(*s*) = − 0.01, ∀*s* ≠ *s*\*). The agent’s goal, therefore, was to find the reward state with as short a path as possible. Whenever the agent found the reward state the agent’s location was reset randomly. We define an “episode” as a single instance of the agent starting in a random location and finding the reward (Fig. 1B). Episodes had a maximum length of simulated time (250 time-steps), and thus, if the agent failed to find the reward state in this time no positive rewards were received for that episode. Each agent experienced multiple episodes for training and evaluation (details below).

### 2.3 Episodic Controller

The central component of an episodic controller is a dictionary of size *N* (consisting of keys, {*k*_1_, …, *k*_*N*_} and values {*v*_1_, …, *v*_*N*_}) for storing events. We refer to this dictionary as the “memory bank”. Each stored event consisted of a state activity vector (which was used as a key for the memory), and an array of returns observed following the selection of one of the four actions (which was the value of the memory). Effectively, the episodic controller stores memories of returns achieved for specific actions taken in past states, and then generates a policy based on these memorized returns. Notably, this means that the episodic controller is not a standard “model-free” reinforcement learning agent, as it does not use a parametric estimator of value. Instead, it uses non-parametric, one-shot memories of experienced returns stored in the memory bank to determine its policies (Lengyel and Dayan, 2007; Pritzel et al., 2017; Blundell et al., 2016).

#### 2.3.1 Storage

Events were logged in the memory bank at the conclusion of each episode. Specifically, the returns for the episode were calculated from reward information stored in transition buffers. Tuples of state, action, and return (*s*_*t*_, *a*_*t*_, *G*_*t*_) for each time-step *t* in the episode were written to the memory bank, i.e. if writing to memory index *i* the key and value were set to *k*_*i*_ = *s*_*t*_ and *v*_*i*_[*a*_*t*_] = *g*_*t*_ (Fig. 2A). For each event to be stored, if the current state was not present in the memory bank (i.e. if *s*_*t*_ ≠ *k*_*i*_ ∀ *i*), a new value array was initialized and the return information was added at the index corresponding to the action selected. If the state was already present in memory, the value array was updated with the most recent return value observed for the given action. Return values were timestamped by the last time that the dictionary entry for that state was updated. This timestamp information was used to determine which entry in memory was least recently updated and would be forgotten (see below).

**Figure 2.**
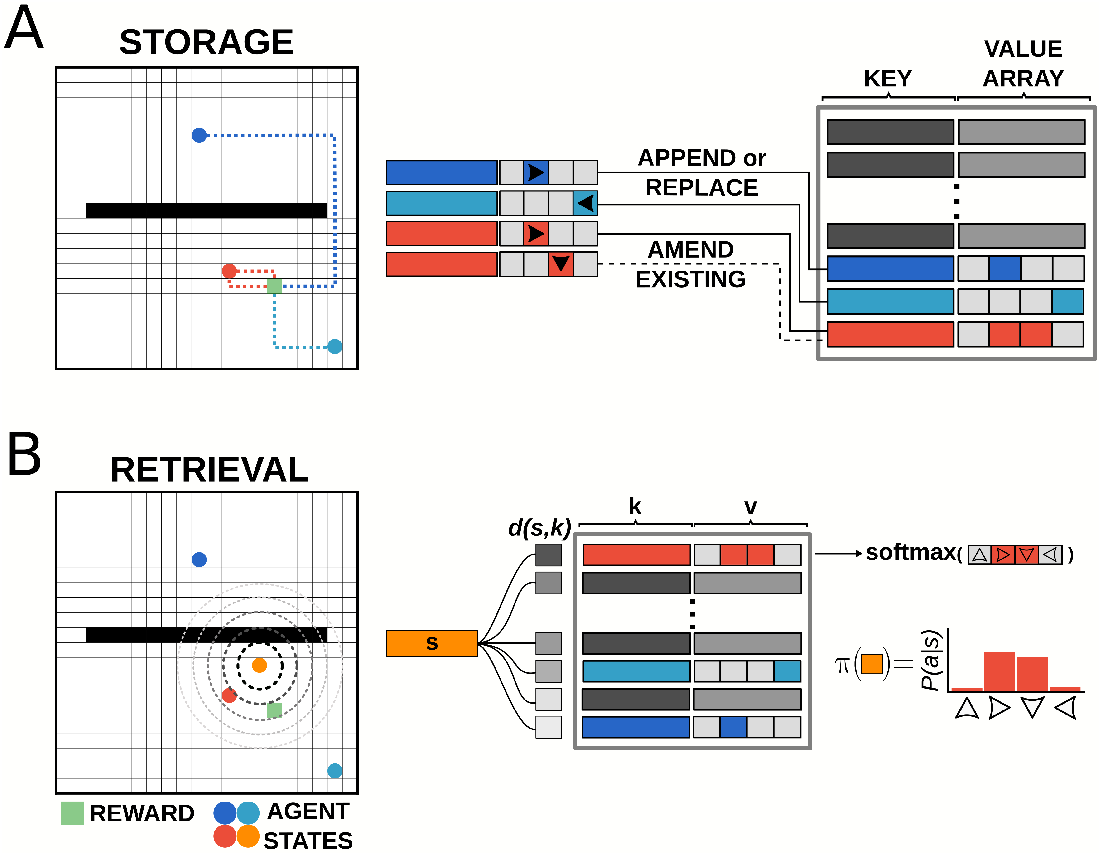
Illustration of episodic control storage and retrieval. (A) Schematic diagram of storage in the episodic controller. The memory bank is a dictionary of key-value pairs, with state activity vectors as keys pointing to value arrays which log returns observed from taking a particular action in that state. Events are written to the memory bank at the end of each episode. Events can be added to memory up to a predetermined number of unique keys (states). If the current state does not exist in memory, a new value array is appended to record the action-return information. If the current state does exist in memory, the value array is amended/replaced with the most recent action-return information. (B) Schematic diagram of retrieval in the episodic controller. Items are retrieved from memory at each time-step of an episode. Pairwise distance between the activity vector for the current state *s* and all state activity keys *k* in memory is computed, and the entry with the smallest distance *d*(*s, k*) is used to generate a policy from the recorded return values.

#### 2.3.2 Retrieval

For each step in an episode, the episodic controller produced actions by sampling from a policy generated by retrieving past events stored in the memory bank (Fig. 2B). On the first episode of each simulation, the episodic controller used a random walk policy to explore the space, as no items were present in memory (since event logging was done at the end of an episode), and thus no policies could be constructed from previous experiences. Once information was logged to the dictionary, the agent generated a policy for each state by querying available states in memory. Namely, if the agent was in a state *s*_*t*_ = *s*, the recall function measured pairwise Chebyshev distance between the activity vector for state *s* and the state activity vectors present in the list of memory keys ({*k*_1_, …, *k*_*n*_}), and then returned the index, *i* whose key had the smallest distance to the current state. This index was used to retrieve the associated return array, *v*_*i*_, which was used to compute the policy to be followed at that time-step. Specifically, a softmax function across the return values at memory index *i* was used to generate a probability distribution over actions, which was the policy:

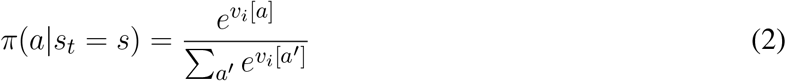

This function maintains the relative magnitudes of the return values present in the associated memory value array. Consequently, actions associated with larger returns would give rise to a greater probability for repeating those actions, while actions associated with smaller returns would be less likely to be selected.

#### 2.3.3 Forgetting

Memory restriction conditions were implemented by limiting the number of entries that could be written to the dictionary. Once this limit was reached, storing a new memory necessitated overwriting a previous memory. In most agents, the entries in memory which were least recently accessed were selected to be overwritten by the new information. Thus, the agents forgot their most remote memories. In random forgetting experiments (see Fig. 8), memories were selected for overwriting by sampling from a uniform distribution over indices in memory.

Memory capacity was set as a percentage of total available (i.e. non-obstacle) states in the environment. For example, when memory was restricted to 75% capacity, we set the size limit of the dictionary to be *N* = 300 for the open field environment, because it had 400 total available states, whereas we set *N* = 273 in the four rooms environment, since it had 365 total available states. The 100% (i.e. unlimited memory) condition was a situation where *N* was set to the total number of available states, and thus, no memory ever had to be overwritten.

### 2.4 State Representations

We compared the performance of an episodic controller under memory restriction conditions using four different representations of state. All representations produced a unique activity vector for each state of the environment (see Fig.3A-B). Unstructured representations contained no relational state information, meaning that activity vectors for states close in graph space shared no common features, and were not close in representation space (Fig. 3C-D). Unstructured representations were either onehot or random state activity vectors. Random activity vectors were produced by drawing samples from the continuous uniform distribution over the half-open interval [0.0,1.0). Onehot vectors were generated by setting a single index to one and zeros in all other positions.

**Figure 3.**
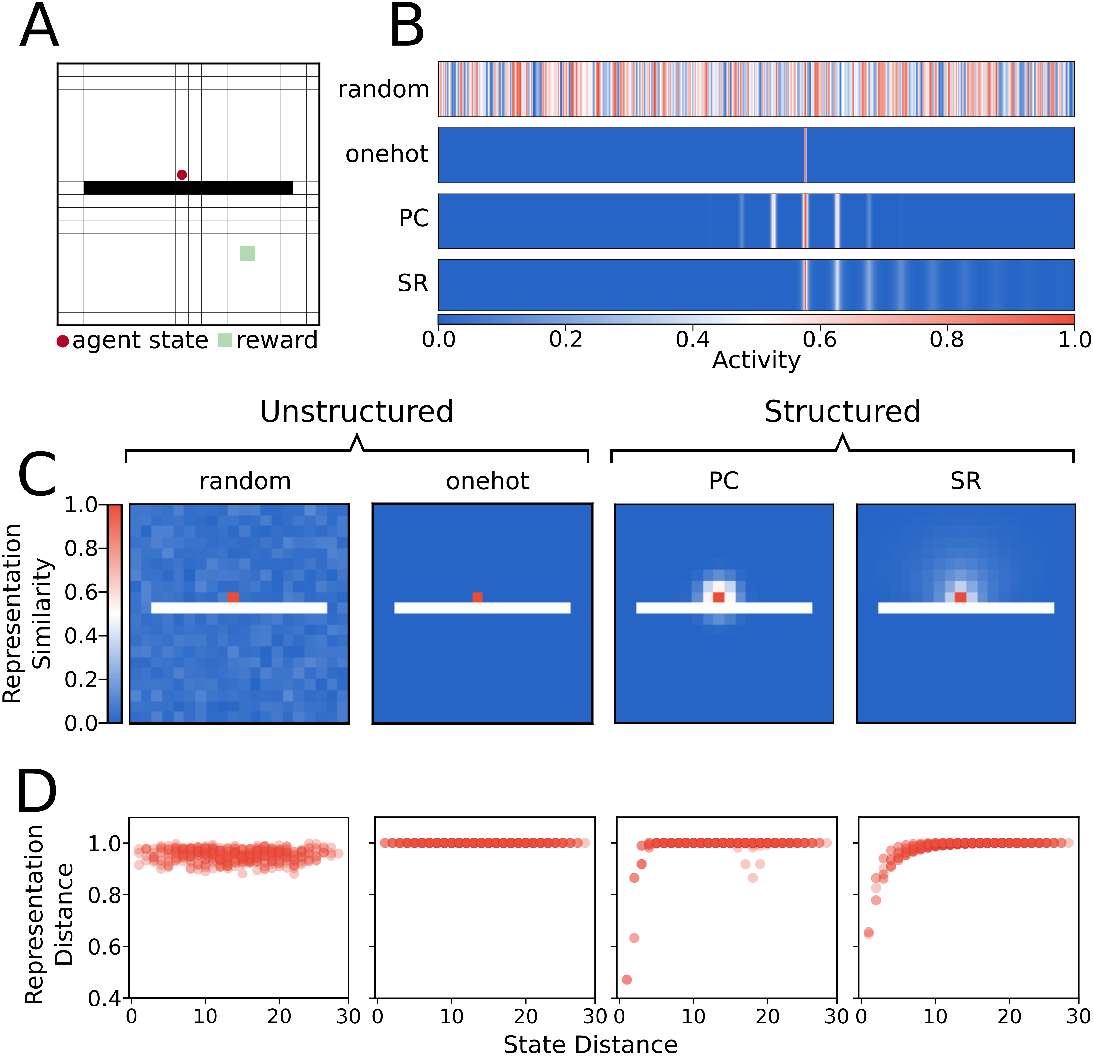
Illustration of state representations used in the simulations. (A) Example state from the separated field environment where the agent is at position (9,10) in the grid. (B) Representations of state (9,10) with random, onehot, place cell (PC), and successor representation (SR) encodings. These state activity vectors are used as keys for the episodic dictionary. (C) Heatmap of Chebyshev (*L*_∞_ norm) distance of state representations for each state and probe state (9,10) for each state encoding. Note that the distance in representation space under random and onehot (unstructured) state encodings has no relationship to the geodesic distance between states in the graph. In contrast, distance in representation space under place cell and successor representation (structured) encodings shows that states nearby on the graph are also nearer in representation space. (D) Distance between representation as a function of geodesic distance between state (9,10) and other states.

In contrast, structured representations encoded a state as a function of its relationship to all other states (Fig 3C-D). Structured representations were either “place cell” or successor representation activity vectors. For place cell representations, each unit of the activity vector was tuned to be most highly activated when the agent state was near its preferred location. The activities of each unit were graded according to how distant the agent’s state was from the preferred location in Euclidean space. Activities were generated by a two dimensional Gaussian function such that when the agent occupied state *s* = (*x, y*), activity of a given unit *i*, with preferred centre (*x*_*i*_, *y*_*i*_) was:

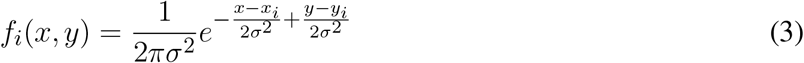

Where *σ* is the size of the place field, here set to be the size of one unit in the grid (1/20 = 0.05).

Successor representation activity vectors described states in terms of the expected future occupancy of successor states. The successor representation for a state *s* is the row of a matrix, *M*, with entries *M* (*s, s*′) given by:

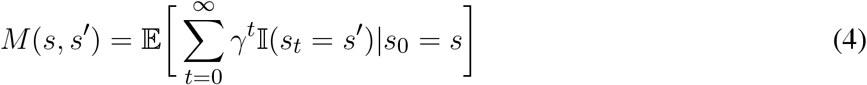

where 𝕀 (*s*_*t*_ = *s*′) is 1 if *s*_*t*_ = *s*′ and 0 otherwise. For a given state transition probability distribution, *P* (*s*_*t*+1_ = *s*′|*s*_*t*_ = *s, a*_*t*_ = *a*), and a given policy, *π*(*a*_*t*_ = *a* |*s*_*t*_ = *s*), the state transition matrix *T* has entries:

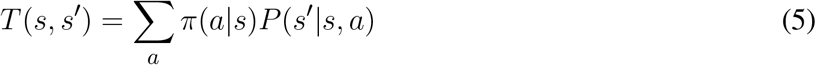

And as such, the successor representation matrix can then be computed as:

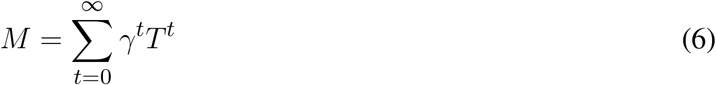

This sum is a geometric series which converges for *γ* < 1, and as such we computed the successor representation matrix analytically by:

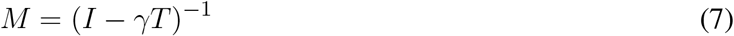

Where *I* is the identity matrix. Notably, we used a random walk policy to generate the successor representations used in these simulations, i.e.*π*(*a*_*t*_ = *a* |*s*_*t*_ = *s*) = *π*(*a*_*t*_ = *a*′|*s*_*t*_ = *s*), ∀*a, a*′. As noted, the successor representation activity vector for each state *s* was the corresponding row of the matrix *M*.

One feature of note is that place cell activity vectors do not respect the existence of boundaries – their activity level is determined only by Euclidean distance between states. By contrast, since the successor representation is computed analytically using the graph adjacency matrix, and edges connecting obstacle states to other states were removed from the graph, the successor representation is sensitive to boundaries (Fig. 3C-D).

#### 2.4.1 Distance Metrics

For states in the graph of the environment, we consider the distance between *s*_1_ and *s*_2_ to be the geodesic distance, i.e. the minimum number of edges which connect these vertices. This geodesic distance respects boundaries, as obstacle states are removed from the graph and as such no edges go into or out of these states. To measure distance in representation space, we use the *L*_∞_ norm, also known as the Chebyshev distance. The distance between two state activity vectors *p* and *q* is given by:

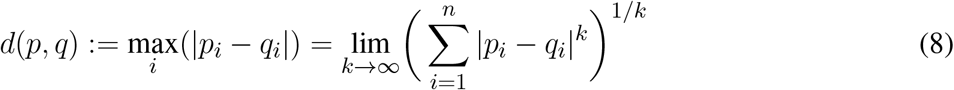

As described above, for a given state *s*, the item in memory used to generate a policy for behaviour was the item at memory index *i* such that *i* = argmin_*i*_*d*(*s, k*_*i*_).

### 2.5 Data Collection

All simulations were run in Python 3.6.8 with functions from NumPy and SciPy libraries. Each simulation was run with a different random seed. Each simulation generated a new instance of an environment class and an episodic controller with an empty memory dictionary. Data was collected over 5000 episodes for each random seed.

### 2.6 Data Analysis

#### 2.6.1 Performance Metrics

To compare between different environments, raw episode reward scores were transformed to percentages of the optimal performance by subtracting the minimum performance score (−2.5) and scaling by the best possible total rewards the agent could achieve, averaged across episodes. To calculate the best possible average cumulative reward across episodes, *R*\*, for each environment we used:

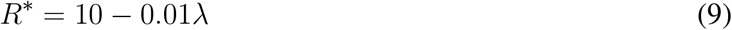

Where *λ* is the mean geodesic distance of all available states to the reward state. This value represents the number of penalization steps an agent with an optimal policy would accrue before reaching the reward location from a randomly chosen starting state of the environment. Thus, the performance of the agent was scaled to be a percentage of the total rewards that an optimal agent would obtain on average. Measures of simulation performance were collected by averaging across the 5000 episodes of a simulation run. Mean performance values for each condition were calculated by taking the mean of simulation-average values over all random seeds. Standard deviation was computed over the simulation-average values. Statistical significance of performance differences between memory restriction conditions was calculated using Welch’s t-test (a two-tailed, unpaired t-test for samples with unequal variances), with a Bonferroni correction used for multiple comparisons. Runs conducted with agents selecting actions from a random-walk policy were used for comparison determining chance levels of cumulative reward.

Successful episodes were those in which the agent reached the rewarded state in fewer than 250 steps. If the agent did not reach the reward state within 250 steps, the episode was terminated, and would be counted as a failed episode.

#### 2.6.2 Policy Maps

For each episode, policy maps were generated by querying the memory for each available state in the environment and storing the resulting policy. These policy maps were used to produce both preferred direction plots (Fig. 6C) and policy entropy plots (Fig. 7), discussed in greater detail below.

To visualize the average policy over time, we generated a two dimensional direction preference vector, **z**_*s*_, for a given state *s* by taking the inner product of the policy and the matrix of 2D cardinal direction vectors, i.e.:

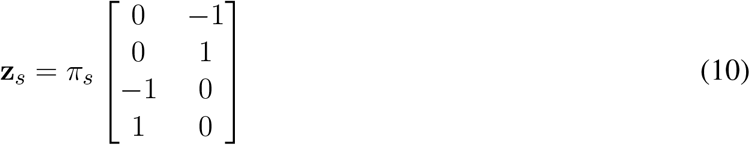

For each state, preferred direction vectors (Fig. 6C) were averaged across the last 400 episodes of the simulation run. This allowed us to average over a large number of episodes while ensuring that for each memory restriction condition, the memory bank had reached its capacity limit.

#### 2.6.3 Trajectories

Example trajectories (Fig. 6A) were collected by reconstructing the episodic memory from saved dictionaries, and sampling actions from episodic policies produced at each state. Trajectories in the unstructured case were taken from an agent using onehot state representations, while trajectories in the structured case were taken from an agent using successor representations (Fig. 3). Three sample trajectories were collected in each memory restriction condition for both structured (successor representation) and unstructured (onehot) state representations from the same three starting locations: (5,5), (5,14), and (14,5). These three starting locations were chosen to visualize paths taken by the agent in the separated field environment such that the agent would have to navigate to the reward from either the same or opposite side of the boundary. All starting states are chosen to be equally distant from the boundaries of the environment such that they have an equal probability of visitation and therefore are equally likely to be present in memory.

To compute average trajectory length, trajectories were sampled from reconstructed episodic memory banks (one per episode, which captured the exact state of the memory at that point in the run). That is, saved dictionaries were used to reconstruct a new episodic controller for each episode. Starting locations were chosen randomly from a uniform distribution over available states, and sample trajectories (n=5) were collected for each episode. The number of steps the agent took (up to a maximum of 250, as in the original runs) was saved for each sample. Trajectory length average (Fig. 6B) was taken over all samples from all episodes together (n=1000). Error is given as standard error of the mean of all samples.

#### 2.6.4 Policy Entropy

The entropy of a policy *π*(*a*|*s*) is the amount of information or surprise inherent in the possible outcomes of sampling from this distribution. The entropy is computed by:

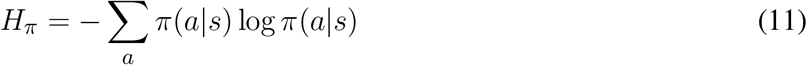

To compute average policy entropy, we first computed policy entropy in each state for each of the last 400 episodes of the simulation run, and then averaged the entropy measure for each state across episodes.

#### 2.6.5 Forgetting Incidence

To measure how the choice of forgetting rule (either forgetting the oldest entry in memory or a random entry) changed which states were more or less likely to be forgotten, we kept a running tally of states discarded from memory for each simulation run. That is, for each state in an environment we maintained a count of the number of times that state was overwritten in the memory bank. We divided these counts by the total number of events of forgetting to get the frequency with which each state was forgotten for a given simulation run. Forgetting frequency arrays were averaged across simulation runs of the same type (structured or unstructured, random or oldest forgetting, *n*=6 for each combination). The relative incidence of forgetting was computed by taking the difference between the average frequency of forgetting under the oldest-state and the random-state forgetting rules. This difference showed how much more likely oldest-state forgetting was to preserve states in memory (forgot less often than random) or to overwrite states in memory (forgot more often than random).

## 3 RESULTS

### 3.1 Moderate forgetting improves performance for structured state representations

We first investigated the effects of memory restriction on performance in four gridworld tasks for agents using either structured or unstructured representations of state information. In all environments, there was no significant difference in mean performance between structured and unstructured representations when the memory capacity was unbounded (100% capacity, see Fig. 4 and Supplementary Fig. S1). In this case, wherein the agent was able to store each state visited, each state representation was a unique alias in memory and retrieval would always return an exact match to the queried state activity vector. Thus, the agent could always generate a policy based on the exact return values observed in that given state.

**Figure 4.**
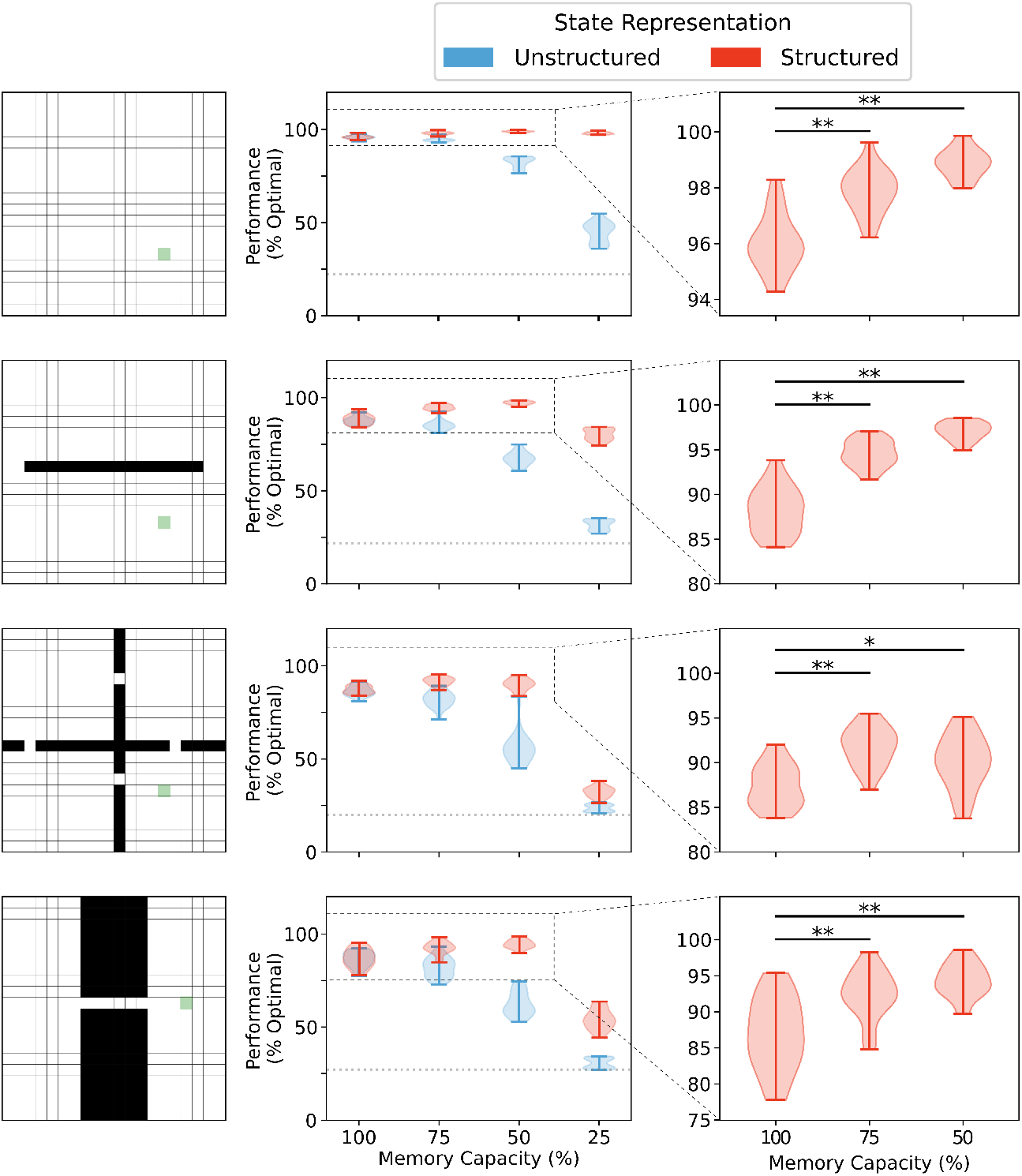
Memory capacity restrictions led to enhanced performance when state representations are structured. No difference in performance was observed between representations when memory capacity is unrestricted (see Supplementary Fig. S1). Structured representations produced better average performance over 5000 episodes when memory capacity was moderately restricted (**, *p* < 1 × 10^−5^; *, *p* < 0.001). Unstructured state representations show a decreased average performance under restriction conditions. At 25% memory capacity (i.e. 75% of all states are forgotten), unstructured state representations performed, on average, little above agents using random walk policies for behaviour (dotted line).

Any restrictions in memory capacity impaired performance of agents using unstructured representations. At the extreme, when only 25% of states encountered were able to be stored in memory, agents using unstructured state representations performed only a little bit better than chance (i.e. rewards collected under random walk policy). In contrast, when memory capacity was moderately restricted, agents using structured representations of state not only didn’t show a reduction in performance, they actually performed better on average than their full-capacity memory counterparts (Fig. 4, *right column*). In general, restricting the size of the memory bank to 60-70% of its full capacity conferred the greatest advantage in most environments (see Supplementary Fig. S2). Importantly, significant restrictions to memory capacity (i.e. only 25% of all states in memory) led to impaired performance regardless of representation type in all environments except the open field. In the open field environment, no significant performance impairment was observed until memory size was restricted to 10% of total capacity (see Supplementary Fig. S2). Thus, for structured mnemonic representations, moderate amounts of forgetting can improve performance of the episodic controller in the foraging task, and the amount of forgetting that can be used is environment-dependent.

### 3.2 Forgetting with structured representations preserves proximity of recalled memories to current state

To better understand the performance of moderate forgetting in agents utilizing structured state representations, we investigated the percentage of episodes in which exact matches to memory were found, and the average distance between representations retrieved from memory and queried state representations (Fig. 5). We found that across all memory conditions successful episodes had a higher percentage of exact matches between stored memory keys and queried states, both for structured and unstructured representations (Fig. 5A). This suggests that the ability to find an exact match in memory is one factor determining performance. Indeed, with forgetting, agents using structured representations maintained a greater proportion of exact match states in memory than agents using unstructured representations (Fig. 5A), and they also performed better in these conditions (Fig. 4). Together, these results imply that with forgetting, agents using unstructured representations experience more failures as a result of an inability to match to queried states, and as the amount of forgetting increases, these agents have fewer exact match to queried states than their counterparts using structured representations.

**Figure 5.**
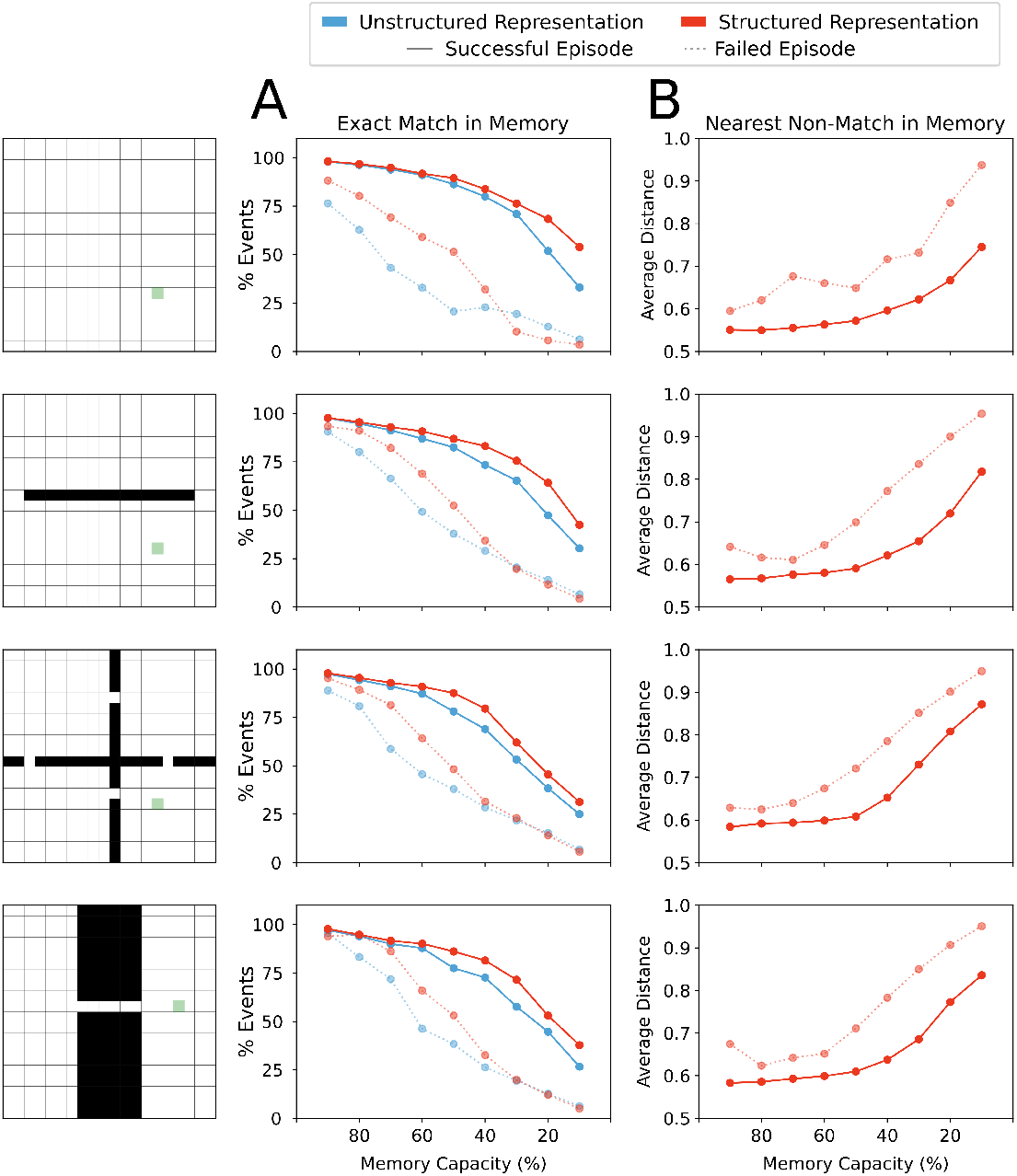
(A) Percentage of events over all episodes in which the agent’s current state in memory was in memory. Agents using either structured or unstructured representations showed a similar incidence of exact match to current state in successful trials when memory capacity restrictions were mild. As memory restrictions were increased, agents using structured state representations maintained a slightly higher incidence of exact matches in memory. By contrast, in episodes in which the agent did not reach the goal state (failed episodes), agents using unstructured state representations showed a lower proportion of exact matches in memory than their counterparts using structured representations. With significant memory restrictions, this difference was eliminated. (B) Structured representations show that the representation retrieved by memory when no exact match was present maintains a close distance to the probe state until memory restrictions become quite severe. On successful trials, agents using structured representations maintained states in memory that were nearer to the current state. In failed episodes, the average nearest state in memory was more distant from the probe state, indicating close neighbours were less often present in memory when the agent was unable to reach the reward state in the allotted time.

**Figure 6.**
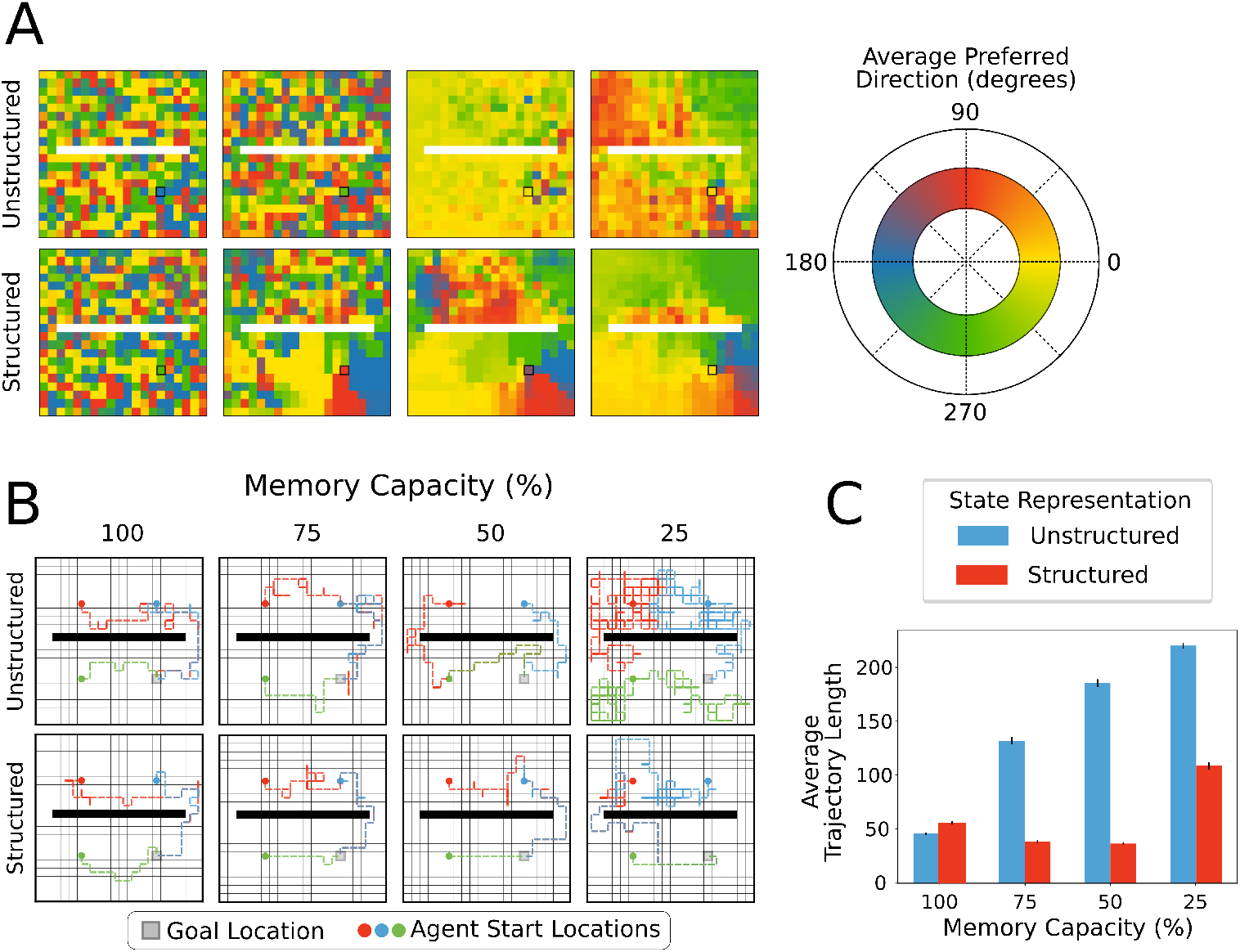
(A) Agents using structured or unstructured state representations displayed similarly low levels of coherence in the policies of neighbouring states when all states could be maintained in memory (i.e. unrestricted). Moderate memory restrictions encouraged neighbouring states to generate similar average preferred directions when structured representations were used, but not when unstructured representations were used. For unstructured representations, decreased memory capacity led to more coherence in neighbouring policies because all states produced the same policies on average, regardless of position relative to the reward location. By contrast, decreased memory capacity led to better policy generalization for structured state representations, especially near the reward location (black square). (B) Example trajectories sampled from episodic controller. Agents using structured representations took more direct paths to the reward location from each of the example starting locations (5,5), (5,14), and (14,5). With more restricted memory capacity, agents using structured representations were able to maintain direct paths to reward, whereas agents using unstructured state representations took more winding paths to reach reward. In 25% memory capacity condition, agents using unstructured representations took paths resembling a random walk. (C) Agents using either type of representation had similar average number of steps per episode (i.e. trajectory length). As memory capacity was restricted, but agents using structured representations reduced average trajectory length, reflecting more direct paths to reward state, while paths taken by agents using unstructured representations became less directed.

**Figure 7.**
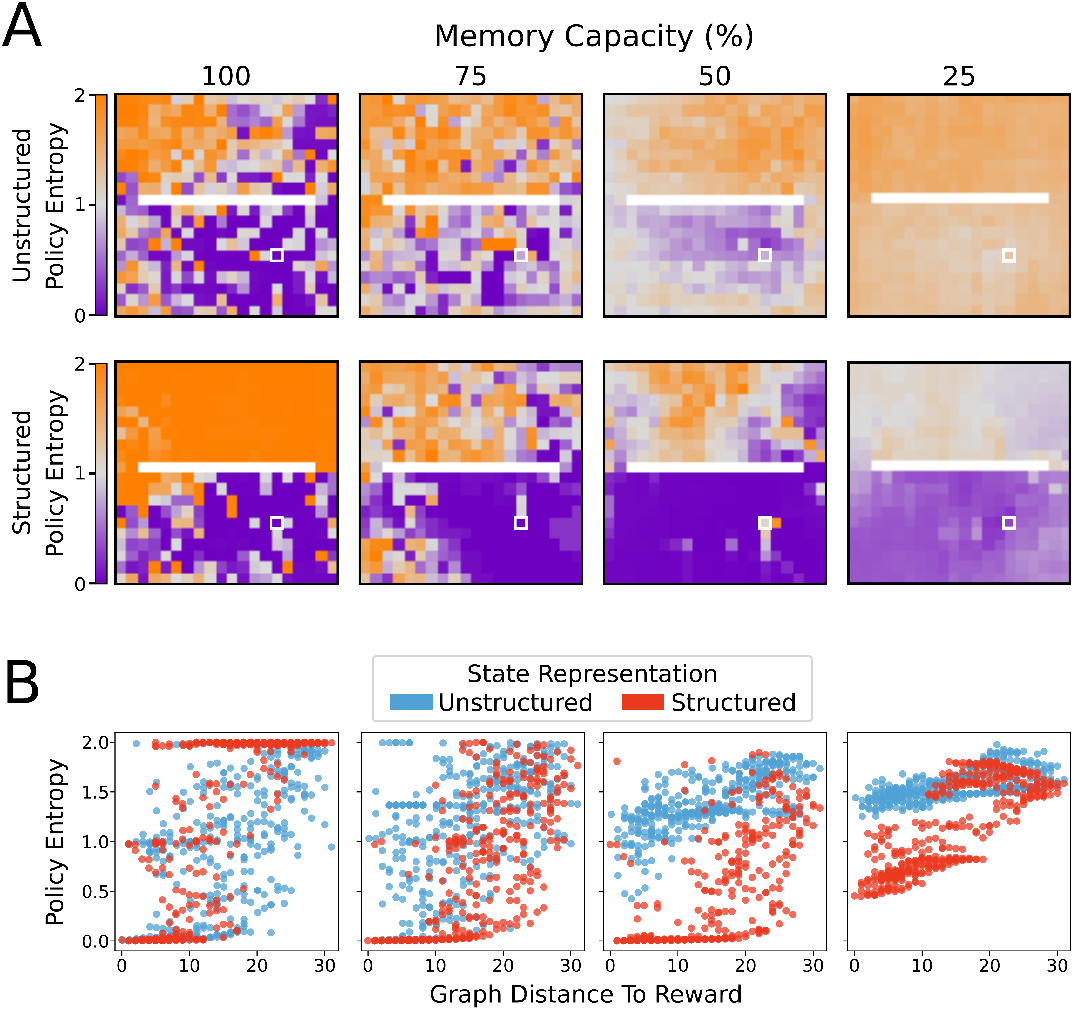
Average policy entropy for each state. (A) Agents using unstructured state representations displayed a greater average policy entropy in all states as memory capacity was restricted. By contrast, limitations on memory capacity promoted lower entropy policies for agents using structured state representations, especially in states more proximal to the reward location (white box). (B) Average policy entropy as a function of geodesic distance from the reward state. Structured representations maintained a lower average policy entropy than agents that used unstructured representations. As memory capacity limitations became stricter, agents using unstructured representations tended to produce more uniformly distributed, higher entropy policies. By contrast, even at most stringent memory restriction conditions, agents using structured representations maintained relatively lower entropy policies at states nearer to the reward location.

**Figure 8.**
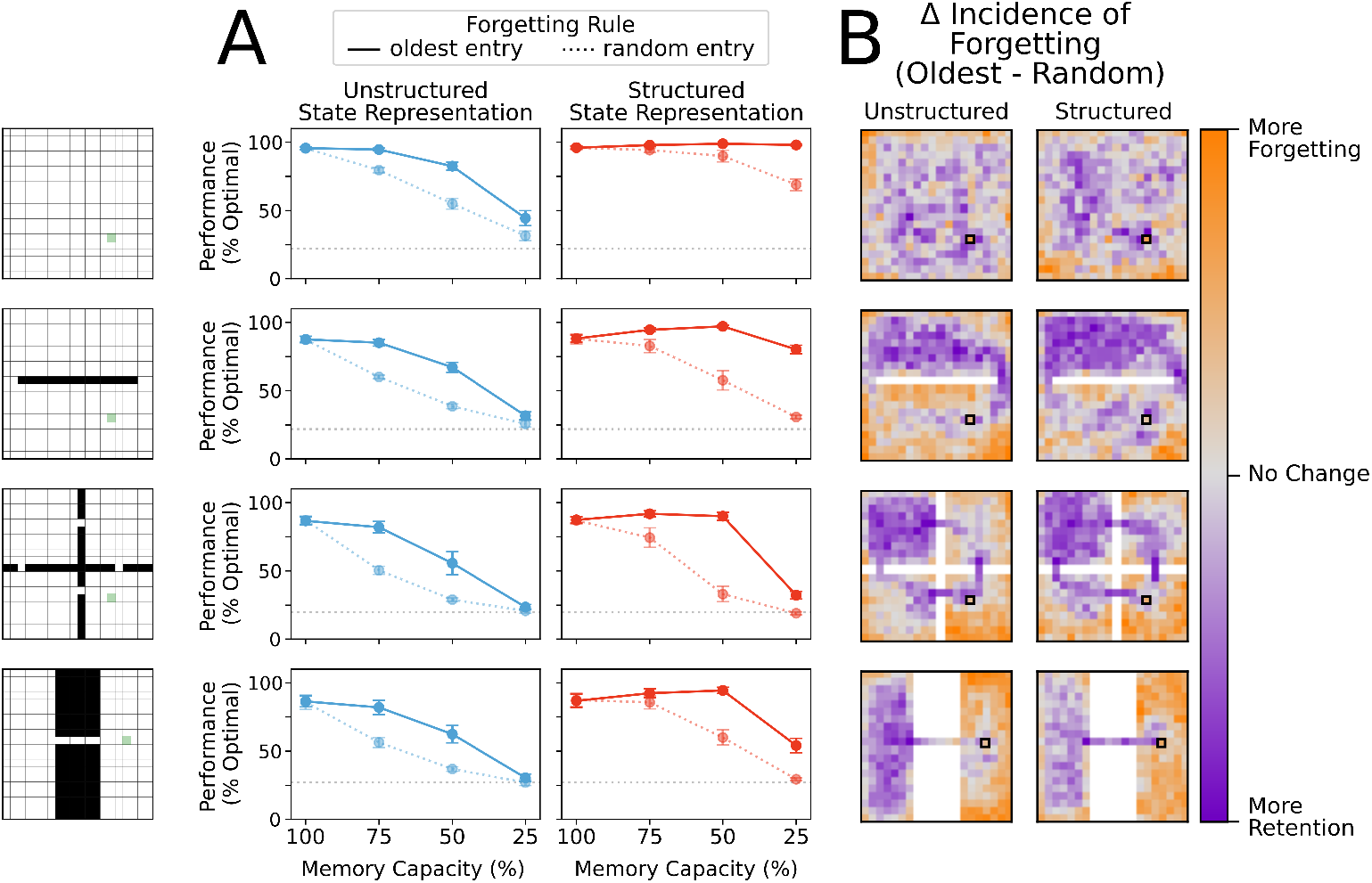
(A) Removal of a random entry from memory impaired performance of agents using both structured and unstructured representations. While agents using structured representations still performed better than agents using unstructured representations, any performance benefit of memory capacity restriction was eliminated. (B) Difference in frequency of forgetting states at 75% memory capacity. Agents using a forgetting rule where random entries in memory were eliminated chose states at the same rate regardless of position. Here we compare the tendency of agents that replace the least recently updated entry in memory to either maintain entries or replace them relative to the random forgetting agents. Agents which replaced the oldest entry in memory showed a greater tendency to preserve bottleneck states and states near the reward location, regardless of state representation. Agents using an oldest-forgetting rule also replaced peripheral states at a higher rate than agents that forgot states randomly.

However, exact matches in memory are clearly not always required for finding the reward, since some successful episodes did not involve exact matches. Therefore, we then explored the average distance to the closest state which was returned by memory when no exact match was present. For unstructured representations, all neighbouring states are equally distant from the probe state (see Fig. 3) and so the nearest distance to a non-match state remains constant regardless of the forgetting condition. In contrast, for structured representations, minimal forgetting produced nearest matches which were relatively close to the queried state in both successful and failed episodes (Fig. 5B). As forgetting increased, the average distance of the nearest match in memory also increased, and we saw a greater increase in average distance on failed trials than for successful trials. This indicated that in trials where the agent was unable to reach the rewarded state within the time limit, it had both fewer exact matches in memory (Fig. 5A) and had on average less similar neighbouring states available in memory from which to generate a policy (Fig. 5B). Moreover, in moderate forgetting conditions (e.g. 90-60% capacity), the distance between states in memory and queried states remained relatively constant, indicating that structured representations allowed the agents to still recall similar states to the queried state under conditions of moderate forgetting. This can explain why moderate forgetting did not have a detrimental impact on episodic control using structured representations.

### 3.3 Moderate forgetting with structured representations promotes policy coherence

Our results on memory retrieval matches helped explain why structured representations did not suffer from performance limitations with moderate forgetting. But why did moderate forgetting produce a slight increase in performance for structured representations? To address this question, we next investigated how forgetting impacted the policies of agents using different state representations. In particular, we wanted to investigate the ways in which policies of neighbouring states agreed with each other or not. In other words, we asked, to what extent does the episodic controller generate a spatially coherent policy under different forgetting conditions? This matters because in the absence of a spatially coherent set of policies the agents may traverse winding paths to the reward, rather than move directly to it. Such a difference could impact performance slightly, given the small negative reward for moving. To visualize this, we computed the preferred direction for each state as the policy-weighted average of the cardinal direction vectors in polar coordinates. This gave an angle which the agent was, on average, likely to move from the given state. This can be thought of as an approximation of the gradient of the policy map.

We found that with unrestricted memory, agents using structured and unstructured representations of state showed similarly low levels of spatial coherence in average preferred direction. Put another way, policies for neighbouring states were not very consistent, and did not tend to recommend similar actions. For agents using unstructured state representations, restricting memory capacity caused neighbouring states to produce more consistent policies, but these policies did not become more likely to lead the agent to the rewarded state. By contrast, restricting memory capacity for agents using structured state representations promoted a high level of policy coherence for neighbouring states, especially for states near the reward location, and these policies were appropriate policies for finding the reward (Fig. 6A). As a result, the average path length for the agents with structured representations, but not unstructured representations, decreased slightly with moderate forgetting, which can likely explain their improved performance (Fig. 6B-C). Another way of understanding this result is that moderate forgetting with structured representations allows the policies to generalize over space more, which can be beneficial in moderation to prevent undue wandering due to episodic noise. This is in-line with previous work showing that reducing forgetting in animals reduces memory generalization (Migues et al., 2016).

### 3.4 Moderate forgetting with structured representations promotes greater certainty in policies

In addition to the impact of forgetting on spatial coherence of the policies, we wondered whether forgetting might also impact performance via the “confidence” of the policies, i.e. the extent to which the agent places a large amount of probability on specific actions. Thus, we measured the average policy entropy for each state for agents using structured or unstructured state representations at different levels of restriction to the memory bank capacity. Here, low entropy policies are those which strongly prefer one action; high entropy policies are those which tend toward the uniform distribution (and are therefore more likely to produce a random action). With unrestricted memory capacity, both agents using structured and unstructured representations were more likely to produce low entropy policies closer to the rewarded state (Fig. 7A, *left column*).

Greater restriction on memory capacity caused agents using unstructured state representations to produce higher entropy policies in more areas of the environment (Fig. 7A, *top row* and Fig. 7B, *blue points*). In contrast, moderate restrictions on memory capacity encouraged agents using structured representations to produce even lower entropy policies near the rewarded state (Fig. 7A, *bottom row* Fig. 7B, *red points*). These results suggest that the agents with structured representations also benefited from moderate forgetting thanks to an increase in their policy confidence near the reward.

### 3.5 Enhanced performance depends on forgetting more remote memories

Finally, we wondered whether moderate forgetting in general was beneficial for performance on structured representations, or whether or design of forgetting the most remote memories was important. Specifically, in forgetting conditions, once the memory bank limit was reached the agents overwrote those memories that were accessed the longest time ago. To determine how important this was for our performance effects, we compared the performance of an agent that replaced the most remote item in memory with an agent that chose random entries for overwriting.

We found that the structured representation performance advantage was eliminated when random items were deleted from memory, and instead, more forgetting always led to performance reductions (Fig. 8A). Moreover, agents using unstructured representations also performed worse when random items were deleted from the memory bank (Fig. 8A). The fact that random forgetting impairs both agents using structured and unstructured state representations suggests that overwriting random states is as likely to delete an important or useful state from memory as it is to delete a relatively uninformative state from memory. Thus, we speculated that agents using random forgetting were more likely to prune states from memory that important for navigation to the reward than agents using a forgetting rule which prioritized removal of remote memories. This speculation was based on the idea that removing remote memories would tend to overwrite states that were more infrequently visited and therefore less important in finding the reward.

To determine whether this was true, we compared the frequency at which a state was forgotten under each of these conditions. Agents using a forgetting rule where random entries in memory were overwritten chose states at the same rate regardless of position. We then visualized the retain/forget-preference of agents using a replace-oldest forgetting rule relative to the replace-random forgetting rule. Agents which replaced the oldest entry in memory showed a greater tendency to preserve bottleneck states and states near the reward location, both for unstructured and structured representations (Fig. 8B). Such states are critical visitation points along many potential trajectories to the reward, and thus, it is natural that their removal impairs performance from multiple starting points. Interestingly, agents that used oldest-forgetting also replaced peripheral states at a higher rate than agents that forgot states randomly, and these states are much less likely to be used in navigating to the reward. These results were consistent across memory restriction conditions (see also Supplementary Fig. S3) These results show that forgetting remote memories can help to preserve critical trajectories in memory while eliminating less useful information for behaviour, in-line with the beneficial forgetting hypothesis (Mosha and Robertson, 2016; Robertson, 2018; Hardt et al., 2013; Richards and Frankland, 2017).

## 4 DISCUSSION

In order to explore the hypothesis that forgetting may sometimes benefit action selection, we investigated the effects of memory restriction on the performance of RL agents using episodic control to navigate a simulated foraging task in four different environments (Fig. 1). The episodic controller stored information about returns observed in each state visited in a given trajectory, which could then be queried in subsequent episodes to generate policies for behaviour (Fig. 2). As a consequence of restricting the maximum number of entries which could be stored in memory, agents were forced to overwrite (i.e. forget) some prior experiences. We measured differences in performance when states were represented with activity vectors which either encoded a state in terms of its position in the more general environmental context (structured representations), or encoded a state as an unique alias unrelated to any other state (unstructured representations, see Fig. 3).

When information for each state could be stored in memory (i.e. no forgetting), there was no difference in performance between agents using structured and unstructured state representations. When all states could be remembered perfectly, there was no need to recall information from nearby states (i.e. state aliasing is trivial), and consequently, there was no need to make use of the relational information contained in structured representations (Fig. 4). Similarly, stringent memory capacity restrictions (only 25% availability) impaired performance regardless of state representation condition (Fig. 4), because such strong restrictions on memory capacity forced the removal of episodic information necessary for navigating through bottleneck states. However, when state representations contained structural information, moderate limits on memory capacity actually enhanced performance. In contrast, this advantage was not conferred on agents using unstructured representations of state (see Fig. 4).

We subsequently explored potential explanations for the performance of structured representations with moderate forgetting. We found that forgetting with structured representations preserved the proximity of recalled memories to the current state. Agents using structured state representations averaged a smaller distance between recalled representations than their counterparts using unstructured state representations, regardless of the degree of forgetting (Fig. 5). In addition, structured state representations promoted similar policies in neighbouring states (Fig. 6A), which then lead to more consistent, and efficient trajectories to the reward (Fig. 6B-C). Unstructured representations, on the other hand, did not promote coherent policies among neighbouring states, so agents still took more meandering trajectories with moderate forgetting. We also observed that agents utilizing structured representations demonstrated greater certainty in the actions they selected (i.e. lower policy entropy), and thus they were more likely to sample “correct” actions (Fig. 7). Finally, we found that these results depended on forgetting remote memories and preserving recent memories, and this seemed to be due to the importance of recent memories for traversing bottleneck states (Fig. 8). Altogether, these results provide theoretical support for the hypothesis that some degree of forgetting can be beneficial because it can help to remove noisy or outdated information, thereby aiding decision making.

This work offers a putative explanation for how active forgetting in biological brains may present advantages over a memory system that can store all events ever experienced. The real world is a complex and dynamic environment in which underlying statistics are not always stationary. Thus, information from prior experience can quickly become obsolete when the statistics from which it was generated have changed. Our results are in-line with the normative hypothesis that active forgetting of experienced episodes can be helpful for behavioural control as it minimizes interference from outdated or noisy information (Richards and Frankland, 2017; Anderson and Hulbert, 2021).

Moreover, our work shows that representations of state with useful semantic content, such as relational information, can leverage similarity between related states to produce useful policies for action selection. A wealth of work in psychology and neuroscience suggests that the hippocampus, the central structure in storage and retrieval of episodic memories, functions to bind together sensory information with relational or contextual information such that it can be used flexibly for inference and generalization (Eichenbaum, 1999; Preston et al., 2004). In accordance with these ideas, we showed that leveraging the relational structure between representations of state can enable generalization from experiences in neighbouring states to produce successful behavioural policies even when a given state is not explicitly available in memory.

To-date, episodic-like memory systems in reinforcement learning tasks have largely bypassed the question of forgetting by allowing memory systems to grow as needed (Lengyel and Dayan, 2007; Pritzel et al., 2017; Blundell et al., 2016; Ritter et al., 2018). We propose that episodic control mechanisms which more faithfully model the transient nature of biological episodic memory can confer additional advantages to RL agents. In particular, including notions of beneficial forgetting and state representations which contain rich semantic information could potentially provide additional performance benefits over agents maintaining unrestricted records.

It should be recognized that this work is limited in its explanatory power because it has only been applied in simple gridworld environments where state information is relatively low dimensional. More complex navigational tasks (i.e. greater number of possible state/action combinations, tasks involving long range dependencies of decisions, etc.) would provide more biologically realistic test beds to apply this conceptualization of beneficial forgetting. Additionally, hippocampal cellular activity involved in representations of episodic information has historically been thought to furnish a cognitive map of space (O’Keefe and Nadel, 1978). Moreover, recent work has demonstrated that the hippocampus appears to encode relational aspects of many non-euclidean (and even non-spatial) tasks (Schapiro et al., 2016; Constantinescu et al., 2016; Aronov et al., 2017; Stachenfeld et al., 2017; Zhou et al., 2019). Thus, in order to make a stronger claim about modeling the effects of forgetting in episodic memory, this work should also be applied in non-navigation tasks.

In addition, the return information used here to generate policies was computed by Monte-Carlo sampling of rewards, which is not the only way animals compute relative value of events in a trajectory (Dolan and Dayan, 2013; Niv, 2009; Toyama et al., 2019). Perhaps the main issue is that Monte Carlo methods require the task structure to be episodic—i.e. trajectories eventually terminate—and they require backwards replay for calculating returns. There is some evidence for such backwards replay in the hippocampus (Wilson and McNaughton, 1994; Ólafsdóttir et al., 2018), but animals are also able to learn in an online fashion in continuous tasks (Niv, 2009; Gershman and Daw, 2017). Indeed, much work in RL and neuroscience has led to the conclusion that animals learn value of states by bootstrapping using temporal differences, with dopaminergic activity of striatal neurons providing a signal of a bootstrapped reward prediction error for learning (Schultz et al., 1997; Montague et al., 1996; Sutton and Barto, 1998). Here, the choice to store Monte-Carlo return values rather than bootstrapped reward prediction errors was done largely to reduce variance in return estimates, but in principle, there is no reason that online value estimates learned with reward prediction errors could not be used.

An additional aspect of forgetting that we did not explore is that the brain tends to prioritize remembering information that is surprising or unique (Brewer, 1988). The forgetting rules presented in this work did not account for violations of expectation of observed rewards or return values. Rather, the primary forgetting rule presented here described information decay only in terms of time elapsed since it was last updated, which could be argued to more closely reflect passive forgetting of information which is not consolidated from short- to longer-term memory stores (Anderson and Hulbert, 2021; Richards and Frankland, 2017).

Finally, this work models behavioural control by episodic memory systems alone. In biological brains, episodic memory is closely interrelated with procedural memory subserved by the striatum (habitual control, roughly analogous to model-free RL) and with semantic memory involving more distributed cortical representations of information (and some work draws parallels between semantic memory and model-based control in RL) Packard and McGaugh (1996); Schultz et al. (1997); Binder and Desai (2011); Gershman and Daw (2017). Experimental work has shown that episodic memories contribute to semantic knowledge by generalization across unique experiences (Sweegers and Talamini, 2014). Moreover, repeated training shifts behavioural control from hippocampally-dependent processing to striatally-dependent processing (Packard and McGaugh, 1996). These findings demonstrate that episodic memory does not function in a vacuum, and that behavioural control in the brain is dependent on a combination of mnemonic processes. Thus, future work could more closely model animal behavioural control joining either model-based or model-free reinforcement learning systems with an episodic control system.

In conclusion, our computational study demonstrates that forgetting can benefit performance of RL agents when representations of state information contain some relational information, and points to potentially fruitful directions for exploring more faithful models of animal behavioural control. Additionally, this work demonstrates that RL systems using episodic control may be enhanced by more faithfully modeling episodic memory as it is understood in psychology and neuroscience, i.e. as a bandwidth limited mechanism to bind sensory and relational information for flexible behavioural control.

## AUTHOR CONTRIBUTIONS

A.Y-C. and B.A.R. planned the study and wrote the paper. A.Y-C. wrote the computer code and generated the data.

## ACKNOWLEDGMENTS

Computations were performed on the Niagara supercomputer at the SciNet HPC Consortium. SciNet is funded by the Canada Foundation for Innovation; the Government of Ontario; Ontario Research Fund - Research Excellence; and the University of Toronto.

## DATA AVAILABILITY STATEMENT

The code to generate and analyze data for this study can be found at github.com/annikc/memory-restricted-EC.

## Supplementary Material

### 1 SUPPLEMENTARY TABLES AND FIGURES

**Figure S1.**
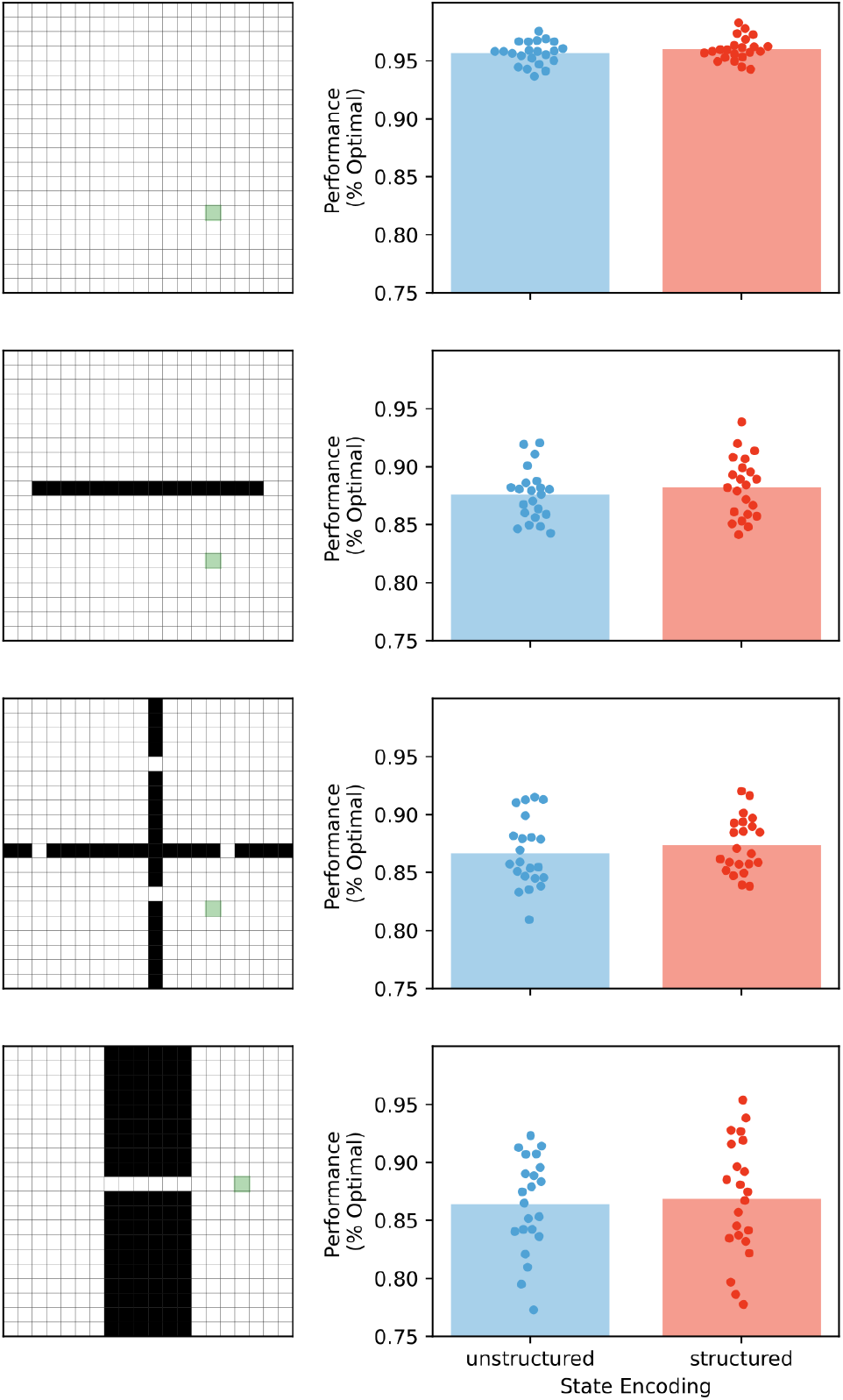
Average performance for episodic control using structured and unstructured representations with unrestricted memory. Performance was computed as the average over 5000 runs for different random seeds (n=22) for each condition. Distribution of average performance was similar between agents using structured and unstructured representations in each of the four gridworld environments. There was no significant difference between the mean performance across random seeds for structured and unstructured representations.

**Figure S2.**
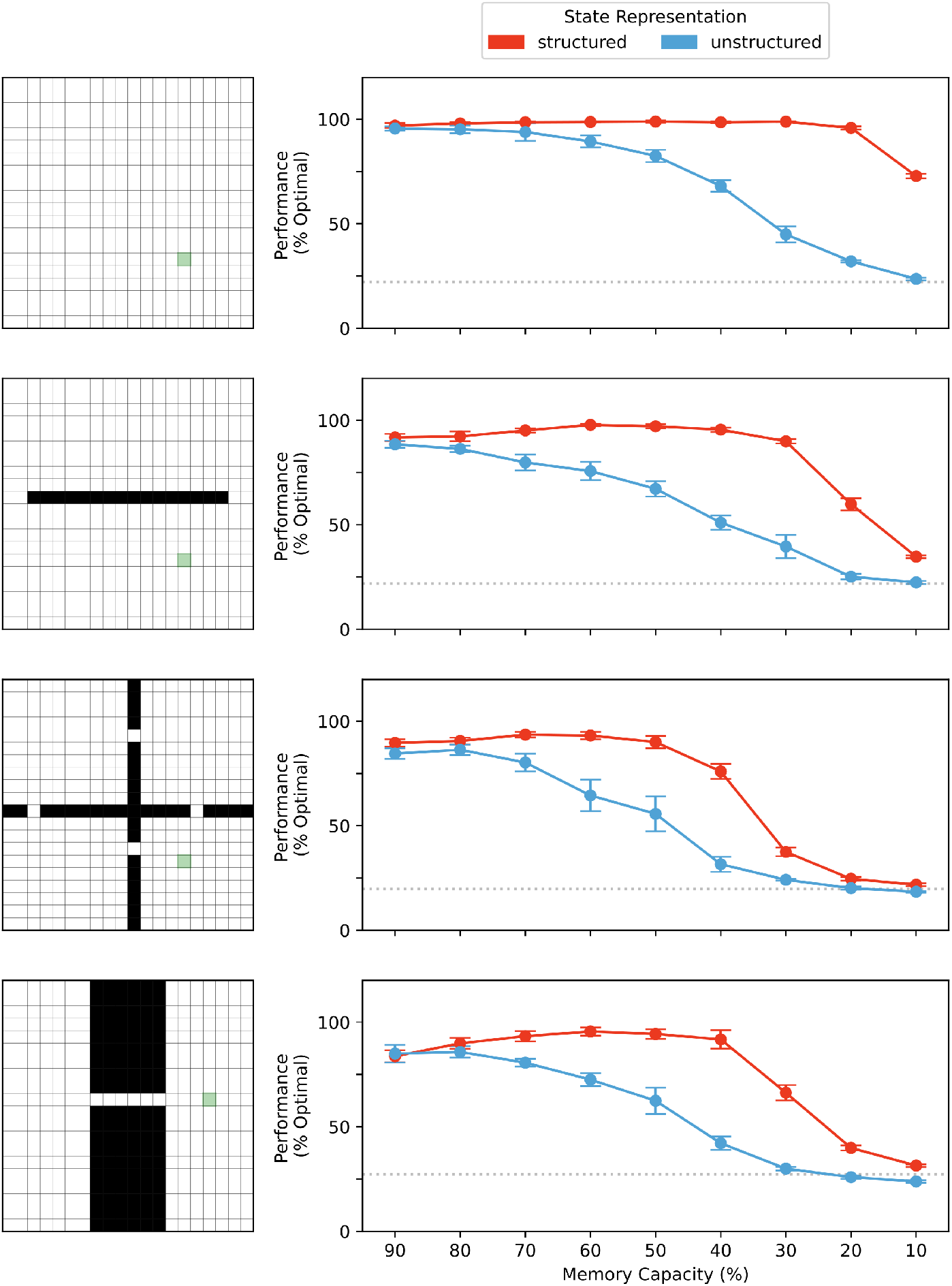
Average performance at memory capacity restrictions in 10% increments for agents using either structured or unstructured representations of state (n=6 for each condition). Data collected and analyzed as in Fig. 4.

**Figure S3.**
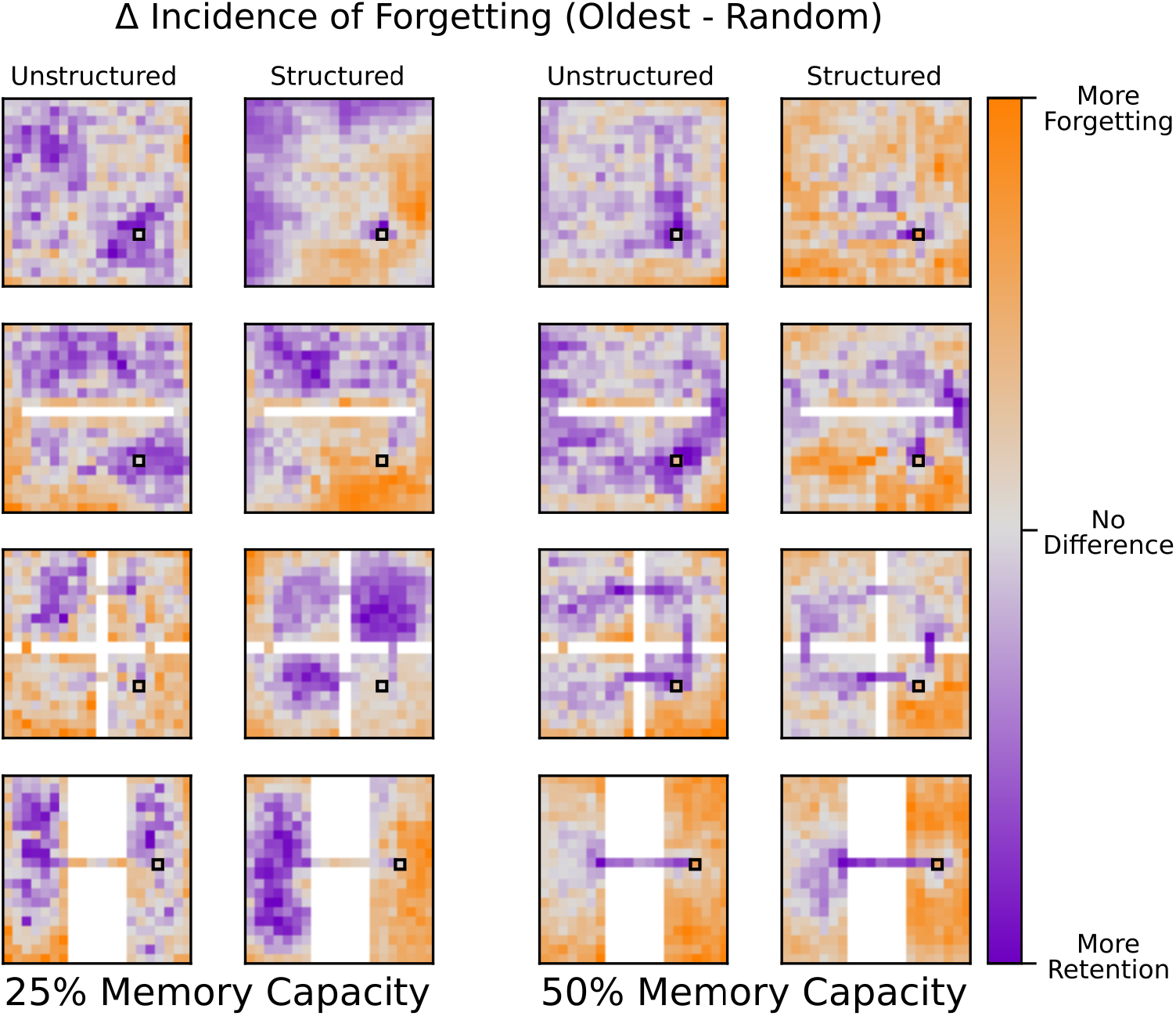
Results as in Fig. 8B for restriction of memory to 50% and 25% capacity. Similarly, the oldest forgetting rule showed a greater propensity for retention of bottleneck states and forgetting of peripheral states over the random forgetting rule condition, except in the case of the open field task for structured representations at 25% memory capacity. In this condition, agents using structured representations and forgetting oldest entries tended to retain memories for states more distal from the reward location at a greater rate than agents using random forgetting.

